# MitoEM 2.0: A Benchmark for Challenging 3D Mitochondria Instance Segmentation from EM Images

**DOI:** 10.1101/2025.11.12.687478

**Authors:** Peng Liu, Boyu Shen, Liyuan Liu, Qiong Wang, Shulin Zhang, Abhishek Bhardwaj, Ignacio Arganda-Carreras, Kedar Narayan, Donglai Wei

**Affiliations:** Boston College; National Cancer Institute; Stanford University; Donostia International Physics Center; University of the Basque Country

## Abstract

We present MitoEM 2.0, a curated data resource for training and evaluating three-dimensional (3D) mitochondria instance segmentation in volume electron microscopy. The collection assembles multiscale vEM datasets (FIB-SEM, SBF-SEM, ssSEM) spanning diverse tissues and species, with expert-verified instance labels emphasizing biologically difficult scenarios, including dense mitochondrial packing, hyperfused networks, and thin filamentous connections with ambiguous boundaries. All releases include native-resolution volumes and standardized processed versions, per-volume metadata (voxel size, modality, tissue, splits), and official train/validation/test partitions to enable reproducible benchmarking. Annotations follow a consistent protocol with quality checks and instance reindexing. Data are provided in NIfTI with nnU-Net–compatible layout, alongside machine-readable split files and checksums. Baseline scripts support common training pipelines and size-stratified evaluation. By consolidating challenging volumes and harmonized labels, MitoEM 2.0 facilitates robust model development and fair comparison across methods while supporting reuse in bioimage analysis, algorithm benchmarking, and teaching.

## Background & Summary

Mitochondria are essential organelles that regulate cellular energy and metabolism. Their three-dimensional structure and organization are crucial for understanding cell function and disease. Volume electron microscopy (vEM) now enables high-resolution 3D imaging of mitochondrial ultrastructure, advancing the quantitative analysis of organelle architecture across cell types. However, automated analysis remains limited by current instance segmentation models, which struggle with challenging cases such as morphologically complex mitochondria, hyperfused networks, and tightly packed organelles with ambiguous boundaries. Although existing benchmarks [1, 2, 3] focus on simpler examples that inflate performance metrics, these difficult scenarios are where automated tools could have the greatest impact. In particular, although the previous MitoEM dataset [4] scaled 3D annotations by more than two orders of magnitude over previous efforts, it provided limited coverage of such difficult cases in hypermetabolic tissues where mitochondrial complexity is most pronounced. This gap continues to hinder the development of robust and generalizable models for biomedical discovery. To address these limitations, we introduce **MitoEM 2.0**, a curated benchmark targeting high-difficulty scenarios in 3D mitochondria segmentation (Fig. 1), with the following contributions:

- **Diversity:** MitoEM 2.0 consolidates vEM volumes from hypermetabolic regions across multiple tissues, species, and imaging modalities, emphasizing difficult mitochondrial phenotypes such as hyperfused networks, slender filamentous connections, dense packing, and ultrastructurally ambiguous boundaries. This provides broad coverage of failure modes.
- **Standardization:** All datasets include native-resolution and standardized processed volumes, consistent instance annotations, machine-readable metadata, and official train/validation/test splits. Uniform file formats, voxel specifications, and evaluation metrics enable reproducible and comparable assessment across datasets.
- **Dataset difficulty quantification:** We introduce two quantitative metrics, morphological complexity and spatial proximity, to quantify instance-level. These metrics expose coverage gaps in existing resources and provide an interpretable structure for organizing datasets within the broader mitochondria segmentation landscape.

**Figure 1.**
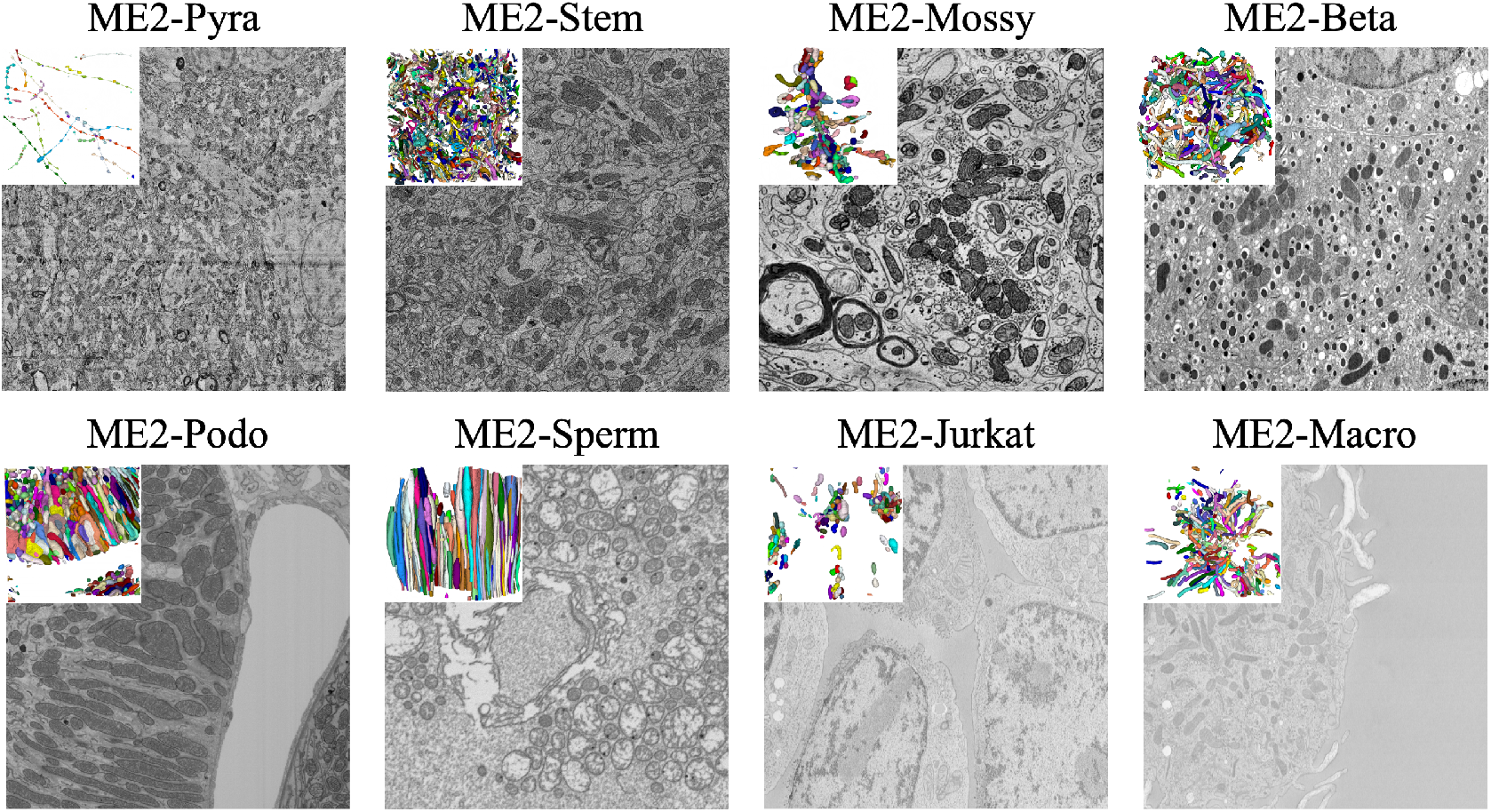
Overview of MitoEM 2.0. MitoEM 2.0 is a collection of 3D volumetric EM datasets with instance-level mitochondrial annotations. It is designed to be diverse, standardized, and difficulty-aware, consolidating challenging volumes from multiple tissues, species, and imaging modalities.

## Methods

### Dataset Curation

The MitoEM 2.0 dataset catalog is organized around biological systems and cell types that pose known segmentation challenges arising from: (1) high mitochondrial density (crowding and small inter-mitochondrial spacing), (2) morphological complexity (branching, fenestrations, irregular surfaces), and (3) restrictive spatial context, including tight packing, confinement in narrow neurites or myofibrillar spaces, and close apposition to membranes/organelles. Our design prioritizes regions where mitochondria exhibit extreme morphological variability and where automated segmentation algorithms commonly fail, ensuring the benchmark captures the full spectrum of real-world segmentation difficulty.

#### Dataset Standardization

All volumes were acquired using volume electron microscopy (EM) modalities, including serial block-face scanning EM (SBF-SEM) and focused ion beam scanning EM (FIB-SEM), across multiple biological specimens. For near-isotropic FIB-SEM data, we downsampled the image volumes to 16nm with linear interpolation and anti-aliasing. For anisotropic ssSEM data, we preserved the original 8×8×30 nm. All datasets include metadata describing the species, tissue type, imaging modality, and voxel resolution.

#### Annotation Procedure

Mitochondria were annotated using a two-step semi-automated pipeline. First, initial segmentations were generated using a fine-tuned 3D U-Net[5] trained on a subset of the data. Next, expert annotators proofread the output using the VAST Lite and Neuroglancer tools, manually correcting under-segmentation, over-segmentation, and merge errors. All mitochondria with a minimum volume of 500 voxels were included. Each mitochondrion was assigned a unique instance label and verified across contiguous slices to ensure topological consistency. To ensure annotation quality, a subset of volumes was independently reviewed by multiple annotators. Discrepancies were resolved through consensus or exclusion. In total, over 8,000 mitochondria instances were labeled across 28 volumes.

### Dataset Details

This section describes each of the eight datasets in MitoEM 2.0, summarizing their biological sources, imaging modalities, and annotation scales. The benchmark integrates volumetric EM data from FIB-SEM, ssSEM, and SBF-SEM across multiple organisms and tissues, capturing wide variation in mitochondrial density, morphology, and resolution. Table 1 provides a unified overview of all datasets. Each entry includes standardized metadata—species, organ, cell type, modality, voxel size, volume dimensions, and train/val/test splits. The catalog spans human, mouse, and Drosophila samples with voxel sizes from 8×8×30 nm to 16×16×16 nm and volumes from 1024×1024×256 to 4096×4096×500, enabling systematic cross-dataset benchmarking under diverse biological and imaging conditions. The dataset details are summarized in Table 1.

**Table 1:** Proposed MitoEM 2.0 benchmark with challenging mitochondria segmentation datasets based on biologically complex regions with dense or morphologically diverse mitochondrial populations. The *Split* column reports the approximate train/validation/test ratio in terms of annotated voxel volume. All FIB-SEM datasets have a resolution of 16 *×* 16*×* 16 nm, whereas ssSEM and SBF-SEM datasets have 8 *×* 8 *×* 30 nm resolution. Existing annotations were refined or converted from semantic masks (^∗^), and new ones were created (^†^).

| Name | Organism | Tissue | Cell Type | Modality | Avg Shape | Voxels | Split | #Mito | EFI/DCI | Ref. |
| --- | --- | --- | --- | --- | --- | --- | --- | --- | --- | --- |
| ME2-Beta | Mouse | Pancreas | β-cell | FIB-SEM | (874,669,979) | 3961M | 3/1/3 | 1935 | 0.02/2.68 | [6]<sup>†</sup> |
| ME2-Jurkat | Human | Cell line | Jurkat | FIB-SEM | (1024,1024,256) | 805M | 1/1/1 | 372 | 0.00/1.62 | [7]* |
| ME2-Macro | Human | Cell line | Macrophages | FIB-SEM | (1024,1024,256) | 805M | 1/1/1 | 497 | 0.01/1.38 | [7]* |
| ME2-Mossy | Mouse | Cerebellum | Mossy Fiber | ssSEM | (1024,1024,500) | 3333M | 3/1/3 | 1099 | 0.10/4.13 | [8] |
| ME2-Pyra | Human | Cerebrum | Pyramidal | ssSEM | (4096,4096,500) | 8388M | 3/1/11 | 45 | 0.96/0.53 | [4] |
| ME2-Podo | Mouse | Kidney | Podocyte | FIB-SEM | (1024,1024,256) | 805M | 1/1/1 | 1333 | 0.00/2.43 | [9]<sup>†</sup> |
| ME2-Sperm | *Drosophila* | Gonad | Spermatocyte | FIB-SEM | (1024,1024,341) | 1073M | 3/1/4 | 775 | 0.01/0.96 | [10]<sup>†</sup> |
| ME2-Stem | Mouse | Brainstem | Mix | SBF-SEM | (1024,1024,100) | 300M | 1/1/1 | 2058 | 0.09/0.80 | [11] |

#### ME2-Beta

(Müller *et al*., 2020 [6]): Mouse pancreatic *β*-cells imaged by FIB-SEM (16×16×16 nm), comprising seven 874×669×979-voxel volumes with 1,935 annotated mitochondria. This dataset exhibits extremely high mitochondrial density and complex morphologies typical of insulin-secreting cells.

#### ME2-Jurkat

(Xu *et al*., 2021 [7]): Human Jurkat T-cells imaged by FIB-SEM (16×16×16 nm), consisting of three 1024×1024×256-voxel volumes with 372 annotated mitochondria, representing mitochondrial organization in immortalized human lymphocytes used in immunological studies.

#### ME2-Macro

(Xu *et al*., 2021 [7]): Human macrophage cell lines imaged by FIB-SEM (16×16×16 nm), comprising three 1024×1024×256-voxel volumes with 497 annotated mitochondria. The dataset captures interconnected mitochondrial networks that support phagocytosis and inflammatory responses.

#### ME2-Mossy

(Han *et al*., 2024 [8]): Mouse cerebellar mossy-fiber boutons imaged by ssSEM (8×8×30 nm), including five 1024×1024×500-voxel volumes with 1,099 annotated mitochondria. It highlights densely packed presynaptic mitochondria supporting high-energy synaptic transmission.

#### ME2-Pyra

(Wei *et al*., 2020 [4]): Human cortical pyramidal neurons imaged by ssSEM (8×8×30 nm). This subset is derived from the original MitoEM dataset by selecting the 17 largest dendritic trees within a 4096×4096×500-voxel volume and extracting all mitochondria contained in them. Despite the modest instance count, the mitochondria span diverse shapes and sizes and are arranged along long, branching dendrites, making this subset particularly challenging.

#### ME2-Podo

(CellMap Consortium, 2024 [9]): Mouse kidney podocytes imaged by FIB-SEM (16×16×16 nm), including two 1024×1024×256-voxel volumes with 1,333 annotated mitochondria. The dataset depicts intricate mitochondrial networks within the specialized filtration structures of glomerular podocytes.

#### ME2-Sperm

(Kunduri *et al*., 2022 [10]): *Drosophila* spermatocytes imaged by FIB-SEM (16×16×16 nm), comprising two 1024×1024×512-voxel volumes with 775 annotated mitochondria. It illustrates distinct mitochondrial arrangements and morphogenetic transitions during spermatogenesis.

#### ME2-Stem

(Jiang *et al*., 2025 [11]): Mouse brainstem tissue imaged by SBF-SEM (8×8×30 nm), comprising three 1000×1000×100-voxel volumes with 2,058 annotated mitochondria. This dataset features mixed neural and glial populations and moderate mitochondrial densities representative of brainstem tissue.

### Dataset Difficulty Quantification

To systematically assess the difficulty of mitochondria segmentation across diverse datasets, we introduce two complementary per-instance metrics: the *Dilation Collision Index (DCI)* and the *Erosion Fragility Index (EFI)*. Together these quantify whether an instance is likely to contact neighbors under small expansions (DCI) and whether its topology is fragile under small contractions (EFI).

- *Dilation Collision Index (DCI)*: For each mitochondrion instance *M*_*i*_ we apply a small morphological dilation with kernel 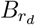 (radius *r*_*d*_) and record the number of distinct other instances that the dilated region touches. In implementation, we use a 3× 3 ×3 structuring element (i.e., *r*_*d*_ = 1 voxel, full 26-neighborhood connectivity). Formally,

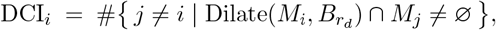

i.e. the cardinality of the set of other instance labels overlapping the dilated mask. A larger DCI indicates that a small boundary expansion would cause contact with many neighbors, i.e., higher likelihood of false merges or boundary ambiguity. Practical computation should ensure unique-label counting (no double counts) and choose a dilation kernel that reflects expected boundary uncertainty
- *Erosion Fragility Index (EFI)*: For each instance *M*_*i*_, we perform a mild morphological erosion using a 3 *×* 3 *×* 3 structuring element (*r*_*e*_ = 1, full 26-neighborhood) and count how many connected components remain. Because the eroded object will always contain at least one ( component corresponding to the original instance, we subtract one to quantify only the newly created fragments:

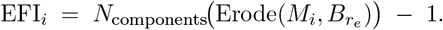

EFI therefore measures topological fragility: higher values indicate that even a slight inward perturbation causes the instance to split into multiple pieces—typical for thin-neck or highly branched mitochondria—and reflect an increased risk of fragmentation or undersegmentation during automated processing.

By projecting mitochondria into the 2D space defined by (DCI, EFI) — we reveal clusters of instances with differing segmentation difficulty: high-DCI/low-EFI objects are densely packed but topologically robust, whereas low-DCI/high-EFI objects are isolated but topologically fragile, for example exhibiting constrictions, thin extensions, or mitochondrial nanotunnels. Many existing benchmarks concentrate in the low-DCI, low-EFI region, leaving a gap in coverage of cases prone to merge- and split-errors. These metrics therefore provide a principled basis to select and evaluate hard instances, and are integral to MitoEM 2.0 dataset design and evaluation framework. A visual comparison of complexity across benchmarks is provided in Fig. 2.

**Figure 2.**
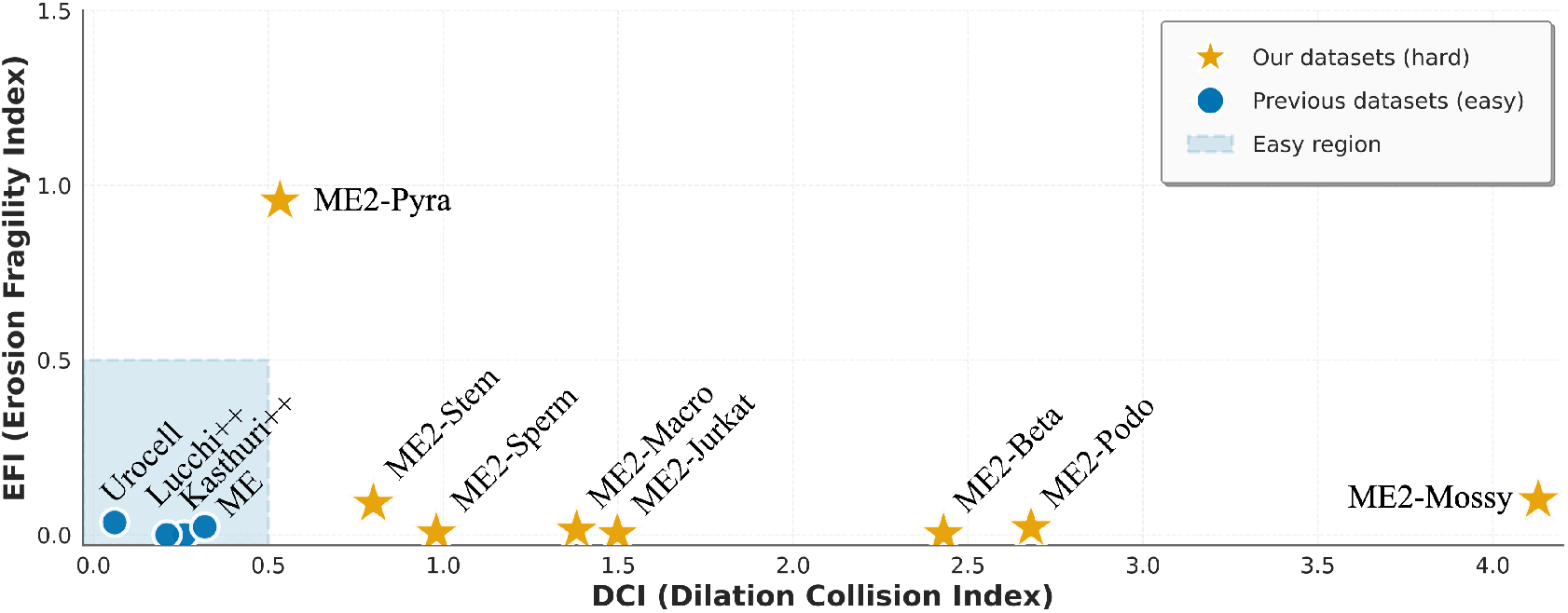
Distribution of test-set difficulty for MitoEM 2.0 and previous benchmarks, measured using the Dilation Collision Index (DCI) and Erosion Fragility Index (EFI). Blue points indicate prior datasets concentrated in a low-DCI/low-EFI “easy region,” while yellow stars denote the eight MitoEM 2.0 datasets, which span higher collision and fragility levels and therefore cover substantially harder segmentation regimes..

### Data Records

MitoEM 2.0 contains eight independently curated datasets, each stored in its own directory (`Dataset001_ME2-Beta` to `Dataset008_ME2-Stem`). All image and label data are provided as 3D NIfTI files (`.nii.gz`) with correct voxel spacing and affine metadata recorded in the NIfTI header. Each dataset follows a consistent, nnU-Net–compatible layout:

1. `imagesTr`/ Training images stored as single-channel `float32` volumes. Filenames follow `me2-<dataset>_trainXX_0000.nii.gz`.
2. labelsTr/ Instance annotations (`uint16`) aligned with `imagesTr`/. Filenames: `me2-<dataset>_trainXX.nii.gz`.
3. `imagesTs`/ Test images following the same convention as training images: `me2-<dataset>_testXX_0000.nii.gz`.
4. `labelsTs`/ Test-set instance annotations: `me2-<dataset>_testXX.nii.gz`.
5. `dataset`.json Machine-readable metadata including voxel spacing, modality, channels, and file lists.
6. `split.json` Official train/validation/test splits used in all baseline evaluations.

A consolidated `metadata.csv` located at the top of the dataset provides biological and imaging attributes for all eight datasets, including system, tissue, cell type, voxel size, modality, and data source. These fields enable filtering and grouping without inspecting individual folders. All volumes are distributed in compressed `gzip` form. Checksums are included to allow users to verify file integrity after download.

### Technical Validation

#### Baseline methods

We evaluate three baseline instance-segmentation methods representative of current practice for mitochondria in EM: (1) MitoNet [3], a Panoptic-DeepLab based generalist model for mitochondria; nnUNet-BC [4] binary mask + contour channel, and (3) micro-SAM [12], a microscopy-adapted version of the Segment Anything Model. We evaluate models under two learning regimes: a *zero-shot (ZS)* setting, where pretrained MitoNet and Micro-SAM models are applied directly without adaptation, and a *from-scratch (Scratch)* or *fine-tuned (FT)* setting, where nnUNet-BC, MitoNet, and Micro-SAM are trained or lightly fine-tuned on each dataset’s designated training split.

##### MitoNet

A Panoptic-DeepLab based 2D generalist for mitochondria (with Empanada/napari tooling for inference and proofreading). Instance masks are produced by center/offset predictions and subsequent grouping/post-processing (optionally with ortho-plane consensus for 3D). For inference on 3D image volumes, MitoNet uses the slice-queue and median-filtering approach with queue lengths adjusted to voxel resolution.

##### nnUNet-BC

A nnU-Net style encoder–decoder that predicts a binary foreground mask together with an auxiliary boundary/contour channel (“BC”). Instance masks are obtained by combining the foreground prediction with the contour channel via watershed/connected-component post-processing to split touching objects. We train the nnUNet-BC model on our training split (same preprocessing and patching as the main experiments), then evaluate on held-out test blocks. Training uses the standard Dice/BCE losses and the same augmentation pipeline described in methods.

##### micro-SAM (SAM)

A microscopy-fine-tuned SAM variant that supports both interactive prompts and automatic instance generation (AMG/AIS). For automatic instance segmentation we run the model’s AIS (Automatic Instance Segmentation) pipeline to produce instances. (1) Zero-shot. use the µSAM pretrained EM organelle weights (no additional training on our dataset) and generate instances using AIS with the default parameter set; this measures out-of-domain generalization and the utility of interactive/foundation-model approaches without dataset specific training. (2) Finetuning. fine-tune EM organelle µSAM model and AIS head on our training split following the same protocol as in methods, then run AIS inference and identical post-processing on the test data.

### Evaluation Metrics

Following the recommendations and retrospective analysis in Franco-Barranco et al. [13], we adopt both instance-level and semantic-level matching accuracy. Both use the same TP/FP/FN framework, but differ in the unit of evaluation: instances versus voxels.

#### Semantic-level accuracy

To quantify voxel-wise agreement between prediction and ground truth, we compute an analogous TP/FP/FN measure at the voxel level. Let *Y*_gt_ and *Y*_pred_ denote binary semantic labels (foreground/background). A voxel is counted as TP (correct foreground), FP (predicted foreground but actually background), or FN (ground-truth foreground missed by the model). Semantic accuracy is defined as

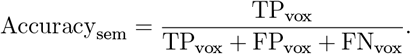

This metric reflects voxel-level foreground/background correctness and is insensitive to object-level splits or merges. Together, the two accuracy measures jointly capture voxel-wise boundary quality and object-level segmentation completeness.

#### Instance-level accuracy

Predicted instances *P* = {*p*_*i*_} are matched one-to-one to ground-truth instances *G* = {*g*_*j*_} using the Hungarian algorithm on an IoU-based cost matrix with a minimum IoU threshold τ = 0.5. After optimal assignment, true positives (TP) are matched pairs with IoU ≥τ,false positives (FP) are unmatched predictions, and false negatives (FN) are unmatched ground truth instances. Instance-level accuracy is defined as

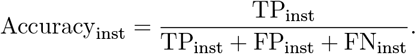

We adopt τ = 0.5 by default and report alternative thresholds where relevant. The cost formulation, tie-breaking, and assignment rules follow the public implementation for reproducibility.

### Benchmark on Each Dataset

We evaluate three representative segmentation pipelines—nnUNet-BC, MitoNet, and micro-SAM—across all eight datasets to assess data usability, quantify dataset difficulty, and provide reference performance for future work. The goal is not to propose new methodology, but to characterize how different architectures behave under the diverse morphological and spatial regimes captured in MitoEM 2.0. The results, summarized in Table 2, reveal consistent and interpretable patterns that reflect intrinsic dataset properties rather than model-specific artifacts. Together, the benchmark highlights five complementary observations, each illustrating a distinct aspect of dataset quality, intrinsic difficulty, or model behavior.

**Table 2:** Per-dataset Accuracy@50. The first half reports **semantic-level** accuracy and the second half reports **instance-level** accuracy (Hungarian matching, IoU threshold *τ* = 0.50). Bold indicates best performance per row for each metric.

| Dataset | Semantic-level Accuracy (↑) | | | | | Instance-level Accuracy (↑) | | | | |
| --- | --- | --- | --- | --- | --- | --- | --- | --- | --- | --- |
|  | nnUNet-BC | MitoNet |  | Micro-SAM |  | nnUNet-BC | MitoNet |  | Micro-SAM |  |
|  | Scratch | ZS | FT | ZS | FT | Scratch | ZS | FT | ZS | FT |
| ME2-Beta | **92.8** | 80.2 | 88.0 | 64.3 | 89.5 | 16.9 | 35.0 | **54.4** | 8.5 | 34.3 |
| ME2-Jurkat | **84.9** | 78.1 | 76.1 | 77.6 | 77.4 | **59.6** | 44.0 | 52.4 | 30.4 | 34.6 |
| ME2-Macro | **89.5** | 74.4 | 87.0 | 78.5 | 87.0 | 30.5 | 30.7 | **43.3** | 16.3 | 40.2 |
| ME2-Mossy | 83.4 | 40.9 | 68.9 | 89.6 | **89.9** | 6.6 | 6.1 | 27.0 | 34.4 | **36.9** |
| ME2-Podo | 90.4 | **91.6** | **91.6** | 87.4 | 91.0 | 27.3 | 35.4 | **58.3** | 25.7 | 39.0 |
| ME2-Pyra | **91.8** | 90.0 | 89.0 | 89.2 | 89.4 | 16.9 | 17.5 | **23.2** | 16.3 | 18.0 |
| ME2-Sperm | 91.6 | 85.9 | 92.0 | 86.9 | **92.7** | 60.8 | 16.8 | **63.7** | 15.6 | 34.6 |
| ME2-Stem | 71.7 | 67.0 | 72.5 | 71.9 | **79.8** | **37.3** | 31.8 | 35.5 | 25.7 | 31.9 |
| Average | 87.0 | 76.0 | 83.1 | 80.7 | **87.1** | 32.0 | 27.2 | **44.7** | 21.6 | 33.7 |

#### Dataset quality: supports stable training

Across all eight datasets, nnUNet-BC, MitoNet, and micro-SAM train reliably without any dataset-specific adjustments. This demonstrates that the annotations, voxel metadata, and standardized directory structure provide clean, well-formed inputs for diverse architectures. Fine-tuning yields clear, consistent improvements for MitoNet and microSAM, confirming that the official training splits adequately capture the underlying morphological variability. Together, these observations indicate that MitoEM 2.0 supports stable, reproducible, and architecture-agnostic training across heterogeneous vEM data.

#### Difficulty quantification: aligns with model performance

Across all models and training regimes, the two lowest instance-level accuracies occur on ME2-Mossy and ME2-Pyra—the datasets with the highest DCI (dense packing) and highest EFI (topological fragility), respectively. This alignment between measured difficulty and observed performance shows that segmentation errors arise from intrinsic biological structure rather than from model-specific failure modes. MitoEM 2.0 therefore provides a benchmark where performance differences are interpretable in terms of organelle density, topology, and continuity, enabling future methods to explicitly target biologically grounded failure modes absent in conventional benchmarks.

#### Segmentation tasks: high semantic accuracy but with low instance accuracy

Semantic-level accuracy is uniformly high across datasets and models, indicating that voxel-wise foreground classification is comparatively straightforward. In contrast, instance-level accuracy remains substantially lower, with frequent merge and split errors, especially in densely packed or topologically fragile regions. This persistent gap demonstrates that the central challenge in MitoEM 2.0 is not detecting mitochondrial pixels but correctly separating individual mitochondria. Instance-level delineation is therefore the dominant bottleneck and the primary target for future algorithmic improvement.

#### Baseline models: exhibits complementary strengths and trade-offs

The three baseline architectures illustrate distinct performance profiles shaped by their design choices. nnUNet-BC, a fully 3D model, achieves the strongest semantic accuracy across several datasets and performs robustly even when trained from scratch. MitoNet achieves the best instance-level accuracy after fine-tuning, benefiting from its center–offset representation and orthogonal-plane consensus. micro-SAM demonstrates strong semantic generalization in both zero-shot and fine-tuned settings but suffers instance-level degradation due to its 2D backbone and slice-wise inconsistencies. Together, these behaviors highlight complementary strengths and illustrate why semantic accuracy alone does not guarantee correct instance reconstruction.

### Usage Notes

All MitoEM 2.0 volumes can be opened using standard biomedical imaging tools such as NiBabel, SimpleITK, ITK-SNAP, Fiji, or Neuroglancer. Users should ensure that their preferred loading library preserves the voxel spacing stored in the NIfTI header, especially when resampling or converting formats.

Instance annotations are stored as `uint16` masks where each mitochondrion is assigned a unique positive integer label (0 denotes background). For semantic-only workflows, these masks can be binarized; instance-aware methods may operate directly on the integer-labeled volumes.

Several datasets, particularly those acquired by ssSEM or SBF-SEM (e.g., ME2-Mossy, ME2-Pyra, ME2-Stem), exhibit anisotropic voxel spacing. Resampling such data to isotropic resolution may introduce interpolation artifacts along the axial dimension. To support diverse analysis pipelines, both native-resolution volumes and standardized processed versions are provided.

Each dataset includes `dataset.json` and `split.json` files defining metadata and official data partitions, enabling direct integration with nnU-Net and ensuring consistent benchmarking. Users performing additional preprocessing (e.g., denoising, registration, downsampling) are encouraged to document these steps to support reproducibility across future studies.

## Data Availability

The MitoEM 2.0 datasets are publicly available on Zenodo at 10.5281/zenodo.17635006.

## Code Availability

All code used to train baseline models, perform inference, and evaluate instance segmentation results is publicly available at https://github.com/luckieucas/MitoEM2.0.

## Author Contributions

D.W. conceived the project and supervised dataset design and curation. P.L., B.S., L.L, Q.W., S.Z and A.B performed the data preprocessing and annotation. P.L. developed the baseline segmentation models and benchmarking pipeline. B.S. and L.L. contributed to annotation tool integration and quality control. P.L. and D.W. conducted technical validation and evaluation. D.W., P.L., K. N., and I.A-C. wrote the manuscript with input from all authors. All authors reviewed and approved the final version of the manuscript.

## Competing Interests

The authors declare no competing interests.

## Acknowledgments

This project has been funded in whole or in part with Federal funds from the National Cancer Institute, National Institutes of Health, under Contract No. 75N91019D00024. The content of this publication does not necessarily reflect the views or policies of the Department of Health and Human Services, nor does mention of trade names, commercial products, or organizations imply endorsement by the U.S. Government. We thank Constantin Pape and Anwai Archit for their constructive feedback and insightful suggestions.

## Funding

This work was supported by the NSF-IIS-2239688. It was also partially supported by grant PID2024-157485NB-I00, funded by the Ministerio de Ciencia, Innovación y Universidades (MICIU/AEI /10.13039/501100011033/) and by FEDER, UE (to I.A-C.).

## Ethics statement

All data analyzed in this work were obtained from previously published, open-access electron microscopy datasets. The generation of these datasets was carried out under the ethical oversight reported in the original publications. Because no new data involving human or animal subjects were collected, the present study did not require additional ethical approval.

## Notes

https://github.com/luckieucas/MitoEM2.0

https://doi.org/10.5281/zenodo.17635006

### Competing Interest Statement

The authors have declared no competing interest.

### Summary of Updates

add funding acknowledge, linke author ORCID

## References

[1] Lucchi P. Márquez-Neila, C. Becker, Y. Li, K. Smith, G. Knott, and P. Fua, “Learning structured models for segmentation of 2-D and 3-D imagery,” IEEE Transactions on Medical Imaging, vol. 34, no. 5, pp. 1096–1110, 2014.

[2] V. Casser, K. Kang, H. Pfister, and D. Haehn, “Fast mitochondria detection for connectomics,” in Medical Imaging with Deep Learning, 2020.

[3] R. Conrad and K. Narayan, “Instance segmentation of mitochondria in electron microscopy images with a generalist deep learning model trained on a diverse dataset,” Cell Systems, vol. 14, no. 1, pp. 58–71, 2023.

[4] D. Wei, Z. Lin, D. Franco-Barranco, N. Wendt, X. Liu, W. Yin, X. Huang, A. Gupta, W.-D. Jang, X. Wang et al., “MitoEM dataset: Large-scale 3D mitochondria instance segmentation from EM images,” in International Conference on Medical Image Computing and Computer-Assisted Intervention (MICCAI). Springer, 2020, pp. 66–76.

[5] F. Isensee, P. F. Jaeger, S. A. Kohl, J. Petersen, and K. H. Maier-Hein, “nnu-net: a self-configuring method for deep learning-based biomedical image segmentation,” Nature methods, vol. 18, no. 2, pp. 203–211, 2021

[6] A. Müller, D. Schmidt, C. S. Xu, S. Pang, J. V. D’Costa, S. Kretschmar, C. Münster, T. Kurth, F. Jug, M. Weigert et al., “3d fib-sem reconstruction of microtubule–organelle interaction in whole primary mouse β cells,” Journal of Cell Biology, vol. 220, no. 2, p. e202010039, 2020.

[7] S. Xu, S. Pang, G. Shtengel, A. Müller, A. T. Ritter, H. K. Hoffman, S.-y. Takemura, Z. Lu, H. A. Pasolli, N. Iyer et al., “An open-access volume electron microscopy atlas of whole cells and tissues,” Nature, vol. 599, no. 7883, pp. 147–151, 2021.

[8] X. Han, X. Lu, P. H. Li, S. Wang, R. Schalek, Y. Meirovitch, Z. Lin, J. Adhinarta, K. D. Murray, L. M. MacNiven et al., “Multiplexed volumetric clem enabled by scfvs provides insights into the cytology of cerebellar cortex,” Nature communications, vol. 15, no. 1, p. 6648, 2024.

[9] C. P. Team, D. Ackerman, M. B. Ahrens, Y. Aso, E. Avetissian, D. Bennett, C. K. E. Bleck, J. Bogovic, M. Bryant, D. Crooks, D. Feliciano, J. Funke, L. Heinrich, H. Hess, M. Innerberger, N. Iyer, M. Kittisopikul, W. Korff, W.-P. Li, W. M. Linehan, Z. Lu, S. Pang, W. Park, K. Pedram, A. Petruncio, A. Post, S. Preibisch, J. Price, W. Qiu, D. Ramirez, J. Rhoades, V. M. S. Ruetten, S. Saalfeld, E. T. Trautman, R. Vorimo, A. Weigel, C. S. Xu, G. Yu, Z. Yu, M. Zouinkhi, and Y. Zubov, “CellMap Segmentation Challenge,” 12 2024. [Online]. Available: https://janelia.figshare.com/articles/online_resource/CellMap_Segmentation_Challenge/28034561

[10] G. Kunduri, S.-H. Le, V. Baena, N. Vijaykrishna, A. Harned, K. Nagashima, D. Blankenberg, I. Yoshihiro, K. Narayan, T. Bamba et al., “Delivery of ceramide phosphoethanolamine lipids to the cleavage furrow through the endocytic pathway is essential for male meiotic cytokinesis,” PLoS Biology, vol. 20, no. 9, p. e3001599, 2022.

[11] Y. Jiang, H. Wang, K. M. Boergens, N. Rzepka, F. Wang, and Y. Hua, “Efficient cell-wide mapping of mitochondria in electron microscopic volumes using webknossos,” Cell Reports Methods, vol. 5, no. 2, 2025.

[12] A. Archit, L. Freckmann, S. Nair, N. Khalid, P. Hilt, V. Rajashekar, M. Freitag, C. Teuber, M. Spitzner, Tapia Contreras et al., “Segment anything for microscopy,” Nature Methods, vol. 22, no. 3, pp. 579– 591, 2025

[13] D. Franco-Barranco, Z. Lin, W.-D. Jang, X. Wang, Q. Shen, W. Yin, Y. Fan, M. Li, C. Chen, Z. Xiong et al., “Current progress and challenges in large-scale 3d mitochondria instance segmentation,” IEEE transactions on medical imaging, vol. 42, no. 12, pp. 3956–3971, 2023.

